# Impacts of sex ratio meiotic drive on genome structure and defense in a stalk-eyed fly

**DOI:** 10.1101/2020.09.23.310227

**Authors:** Josephine A. Reinhardt, Richard H. Baker, Aleksey V. Zimin, Chloe Ladias, Kimberly A. Paczolt, John H. Werren, Cheryl Y. Hayashi, Gerald S. Wilkinson

## Abstract

Some stalk-eyed flies in the genus *Teleopsis* carry selfish genetic elements that induce *sex ratio* meiotic drive (SR) and impact the fitness of male and female carriers. Here, we produced a chromosome-level genome assembly of the stalk-eyed fly, *T. dalmanni*, to elucidate patterns of genomic divergence associated with the presence of drive elements. We find evidence for multiple nested inversions along the *sex ratio* haplotype and widespread differentiation and divergence between X^SR^ and X^SR^ along the entire chromosome. These include a striking X^SR^-specific expansion of an array of partial copies of *JASPer*, a gene necessary for maintenance of euchromatin and regulation of transposable element expression (TEs). In addition, the genome contains tens of thousands of TE insertions and hundreds of transcriptionally and insertionally active TE families. Moreover, we find that several TE families are differentially expressed and/or present at a different copy number in SR male testes, suggesting an association between these two categories of selfish genetic elements in this species. We identify *T. dalmanni* orthologs of genes involved in genome defense via the piRNA pathway, including core members *maelstrom, piwi* and *Argonaute3*, that have diverged in sequence, expression or copy number between the SR and standard (ST) X chromosomes, consistent with altered TE regulation in flies carrying a *sex ratio* X chromosome. Overall, the evidence suggests that this ancient X^SR^ polymorphism has had a variety of impacts on repetitive DNA and its regulation in this species.

## Introduction

The genome was once thought to be little more than a blueprint needed to accomplish biological functions and reproduction of an organism. Yet research in the past few decades has demonstrated that genomes of most organisms are heavily colonized by Selfish Genetic Elements (SGEs) with their own evolutionary interests (Werren, Nur, & Wu 1988; Burt and Trivers 2006; Werren 2011; McLaughlin and Malik 2017). The most ubiquitous category of SGE are transposable elements (TEs). TEs comprise at least 50% of the human genome (International Human Genome Sequencing Consortium 2001), but most are inactive remnants of past invasions. In the model dipteran *Drosophila melanogaster*, TEs make up about 20% of the genome (Hubley et al. 2016), with up to one-third actively producing new insertions (McCullers and Steiniger 2017). Transposable elements may negatively impact hosts in a variety of ways, such as disrupting genes, causing (by their repetitive nature) ectopic recombination (Langley et al. 1988), repressing nearby gene expression (Sienski et al. 2012; Lee 2015; Lee and Karpen 2017), and increasing costs of DNA replication and storage (Badge and Brookfield 1997). Of course, like any type of mutation, TE insertions can occasionally be beneficial (Miller et al. 1997; Sinzelle et al. 2009; Jangam et al. 2017), but the overall negative impact of TEs on organismal fitness is sufficiently strong that hosts have repeatedly evolved diverse genomic defenses against them (Selker 1990; Muckenfuss et al. 2006; Vagin et al. 2006; Aravin et al. 2007; Wolf and Goff 2009). RNAi systems are a primary line of genome defense against TEs in Metazoa. In flies, these defense systems include piwi-associated RNA (piRNA) (Aravin et al. 2007; Zhang et al. 2012; Le Thomas et al. 2013) as well as endogenous small interfering RNA (siRNA) (Kawamura et al. 2008).

Another well-studied category of SGE are meiotic drivers. These elements spread by manipulating gametogenesis in their favor leading to greater than 50% representation of the driver in mature gametes (Lyttle 1991). If drivers are on a sex chromosome, a skew in the sex ratio of offspring will result. Such sex ratio distortion may cause population collapse or even extinction (Hamilton 1967) but can be maintained stably (reviewed in Jaenike 2001; Lindholm et al. 2016) if impacts on carrier fitness counterbalance the advantage of drive to the element (Curtsinger and Feldman 1980). Drive elements can also persist if their action is suppressed (eliminating distortion of the sex ratio); however, enhancers may also evolve, potentially leading to a cyclical arms-race between enhancers and suppressors of drive (Hall 2004). Meiotic drivers are often associated with chromosomal inversions (Jaenike 2001). Among autosomal drivers this occurs because linkage disequilibrium between the driver and its target must be maintained, else distorting chromosomes would drive against themselves (so-called “suicide chromosomes”). Some X-linked drivers are also found in inversions (Wu and Beckenbach 1983; Dyer et al. 2007; Paczolt et al. 2017; Fuller et al. 2020), while others are freely recombining single loci (Montchamp-Moreau et al. 2006). Inversions may be associated with X-linked drivers if the drive requires multiple loci to function. Drive-associated inversions may recombine when homozygous, but depending on their frequency or effects on fitness, recombination may occur rarely, not at all, or instead cause shorter stretches of exchange via gene conversion (Korunes and Noor 2019). A reduction in recombination slows or prevents purging of new deleterious mutations (Muller 1964) and may also reduce nucleotide diversity (Smith and Haigh 1974; Charlesworth et al. 1993). Ultimately, similar to transposable elements, meiotic drive elements can persist long-term in populations despite negative effects on individuals carrying them – although production of excess females has been theorized to contribute to success in interspecific competition (Unckless and Clark 2014; Mackintosh et al. 2021).

Here, we analyze the impacts of selfish genetic elements within the genome of a stalk-eyed fly (*Teleopsis dalmanni*). In this species 10-30% of males possess X-linked elements that prevent proper development of Y-bearing sperm and result in carrier males producing 90% or more daughters (Presgraves et al. 1997). This *sex ratio* (SR) X chromosome has multiple impacts on individual fitness (Wilkinson et al. 2006; Finnegan et al. 2019; Meade et al. 2019) including reduced sexual ornament (eyespan) size in SR males (Wilkinson et al. 1998; Johns et al. 2005; Cotton et al. 2014). The SR X chromosome (X^SR^) appears to have originated approximately 500 Kya (Paczolt et al. 2017), and hundreds of mostly X-linked genes are differentially expressed in the testes of SR males (Reinhardt et al. 2014). X^SR^ contains at least one large chromosomal inversion compared to the standard arrangement (X^ST^) and likely more, as recombination has not been detected in X^SR^ / X^ST^ females (Johns et al. 2005; Paczolt et al. 2017). Recombination occurs between X^SR^ haplotypes in homozygous females but the rate of recombination is about half that in X^ST^ females (Paczolt et al. 2017). Reduced recombination and effective population size has likely contributed to drastically reduced polymorphism on X^SR^ (Christianson et al. 2011).

But how much differentiation has occurred between X^SR^ and X^ST^ in this species? How many inversions are on the X? To what extent have TEs colonized the genome, and are there differences in their expression, copy number, or regulation on X^SR^ chromosomes that might indicate evolutionary interactions with meiotic drive? Can we identify likely candidate genes involved in establishing the meiotic drive phenotype? To answer these questions, we created a chromosomal genome assembly for *T. dalmanni*, annotated transposable elements and genes, and then combined RNAseq, pooled short-read and long-read resequencing data from males exhibiting sex ratio to identify sequence, copy-number and expression differences between the two types of X chromosomes.

## Results

### A chromosome length assembly of the *Teleopsis dalmanni* genome

A primary assembly of the genome of *Teleopsis dalmanni*, a stalk-eyed fly from southeast Asia, was assembled using MaSuRCA from hybrid sequencing data containing long-read and short-read sequences (**Table S1**). After haplotig filtering, the assembly was scaffolded using chromatin conformation information and produced three chromosome-length scaffolds with a total size of 438.2MB, comprising 95.7% of the filtered MaSuRCA assembly. We validated the assembly by comparison to an independently generated linkage map produced using a backcross family from a prior QTL study (Wilkinson et al. 2014) (**Figure S1**). While the maps were largely concordant, a 13.3MB region (61-74MB) is inverted between the assembly and the linkage map, consistent with an inversion difference between the two populations used in the QTL study. BUSCO analysis confirmed the presence of 96.7% of 3,285 conserved dipteran genes (**Table S2**), with 2.0% of BUSCO genes duplicated. Overall, 89.8% of 1-to-1 *Drosophila melanogaster* orthologs are located on the same Muller element in these two Schizophoran fly species (**Figure 1A**). As previously reported (Baker and Wilkinson 2010; Vicoso and Bachtrog 2015), the 97.2 MB *Teleopsis* X chromosome is orthologous to chromosome 2L in *D. melanogaster* (Muller element B). The two autosomes, previously (Baker and Wilkinson 2010) referred to as “C1” and “C2” are similarly sized (176MB and 165MB). C1 consists of Muller D and A (chromosomes 3L and X in *D. melanogaster*), and C2 contains Muller C, F, and E in that order (chromosomes 2R, 4, and 3R in *D. melanogaster*). We also produced a draft assembly of a closely-related cryptic species (Christianson et al. 2005; Paczolt et al. 2017), *Teleopsis dalmanni sp 2 (Td2)* to polarize molecular evolution. This assembly contains 50,545 scaffolds and is less complete (Diptera BUSCO = 90.0%, N50 = 35,545) than the *T. dalmanni sensu stricto* assembly.

**Figure 1.**
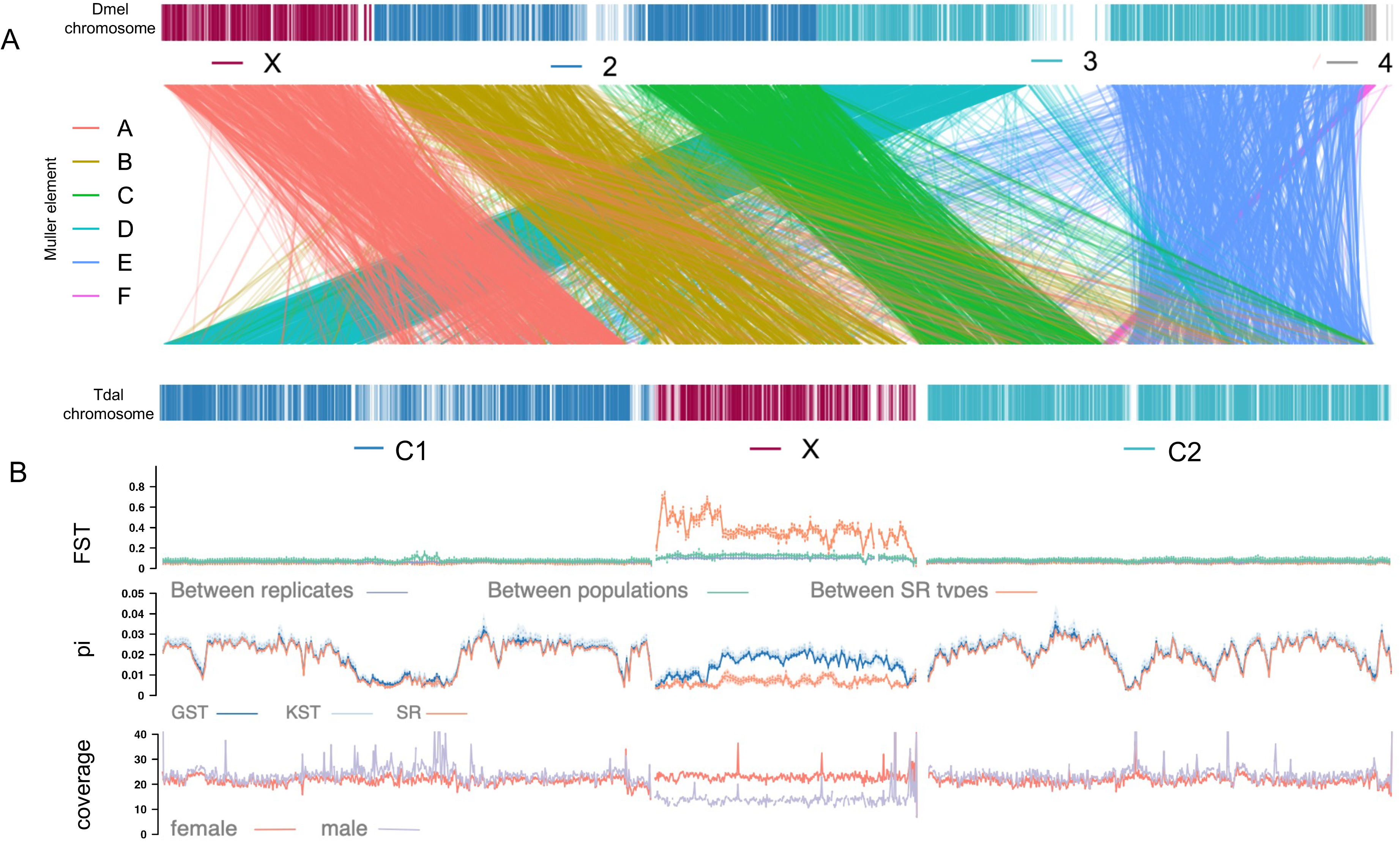
A chromosome length assembly of the stalk-eyed fly genome reveals gene synteny and movement compared to *Drosophila melanogaster*, and unique patterns of sequence variation and differentiation on the X chromosome influenced by meiotic drive. (A) Slopegraph indicating the locations of 7,634 *T. dalmanni – D. melanogaster* 1-to-1 orthologs in each genome. Genes are colored by their Muller element in *D. melanogaster*, with 89.8% found on the same Muller element in both species. As previously reported, the X chromosome in *T. dalmanni* is Muller element B (left arm of chromosome 2 in *D. melanogaster)*. (B) pairwise F_ST_ between pool-seq libraries shows little differentiation between standard pools from the same or different collection sites on any chromosome, but elevated FST across the X between SR and ST pools, whereas Nucleotide diversity (pi) estimated from pool-seq of males with extreme female-biased sex ratios (SR) or with standard sex ratios (ST) from two collection sites is reduced near putative centromeric regions and lower overall on the X chromosome in all libraries. As expected the X chromosome has reduced WGS coverage (reads per bp) in a male library compared to a female library.

### The sex ratio X is diverging from the standard X due to multiple overlapping inversions

We compared patterns of genetic variation on the autosomes and X chromosomes using short-read resequencing data from two pools of sex ratio (SR) and four pools of standard (ST) males derived from two field sites near Kuala Lumpur, Malaysia. As expected, autosomal nucleotide diversity (π) and differentiation (F_ST_) did not differ between pools of SR and ST males from the same collection sites, though there was minor genome-wide differentiation between sites (**Figure 1B**). Also as expected, genomic coverage in females was approximately twice that in males across the X, but not the autosomes (**Figure 1B**). All three chromosomes contain regions with reduced nucleotide diversity in all pools. On the two autosomes, these are near the center of the chromosome, where the Muller elements transition and are presumably centromeric regions (Begun and Aquadro 1992; Begun et al. 2007). Given a similar pattern at the proximal end of the X chromosome (**Figure 1B**) we infer this region contains the centromere of the telocentric X chromosome. Overall, X^ST^ has less nucleotide diversity (0.015670+/− 0.001410 CI) than autosomal sequence from ST (0.021375 +/−0.001432) or SR (0.020134 +/− 0.0000839) male pools. Consistent with previous findings (Christianson et al. 2011), X^SR^ has even lower diversity (0.0057492 +/− 0.001775) (**Figure 1B**). Genetic differentiation (F_ST_) is elevated across the X chromosome between *sex ratio* types (X^SR^ vs X^ST^) relative to differentiation between sites or replicate pools from the same site (**Figure 2**).

**Figure 2.**
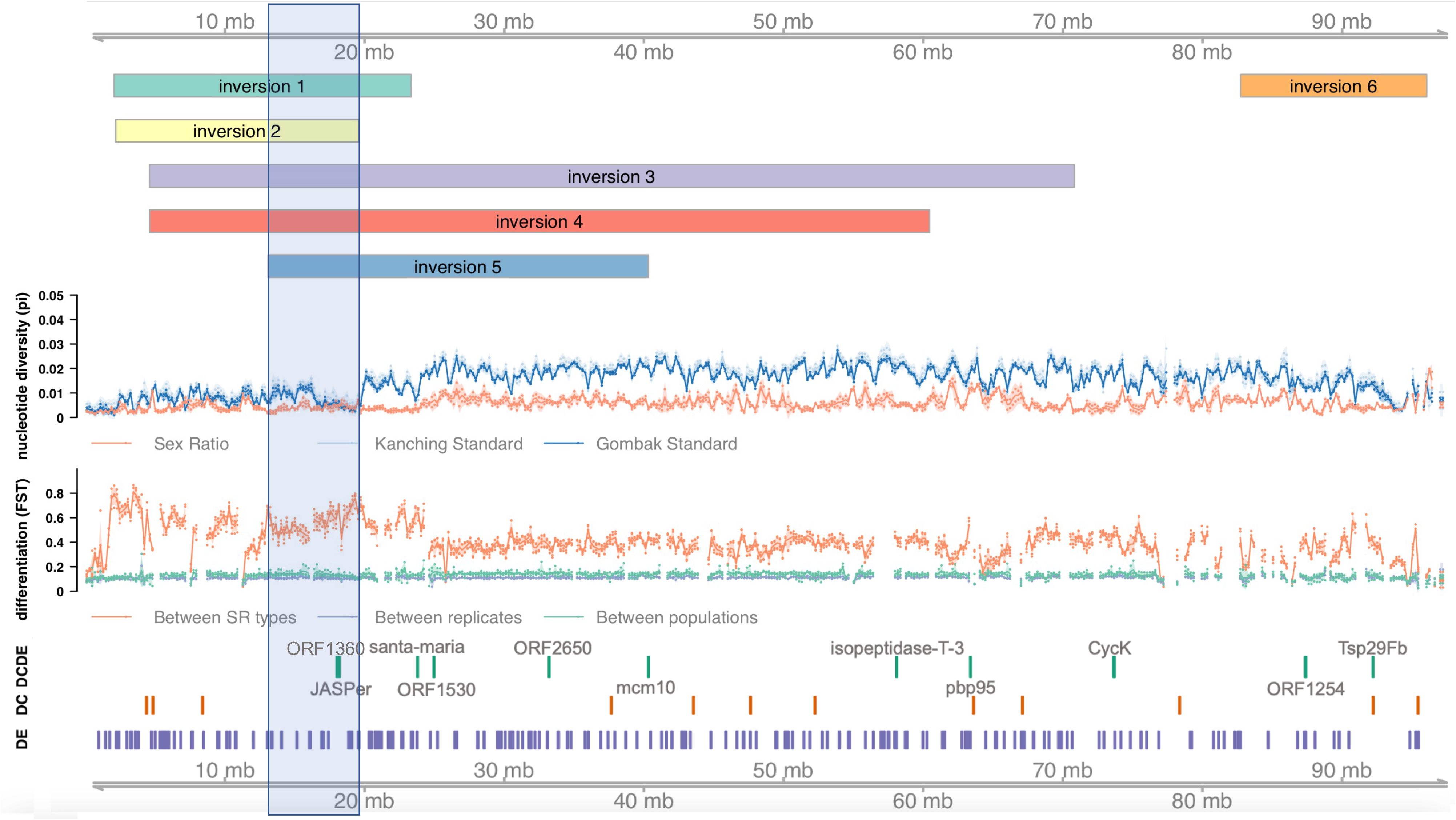
Inversions span the *T. dalmanni* X chromosome, leading to extensive differentiation and divergence in gene expression between SR and ST chromosome haplotypes. Six inversions were identified using PacBio long-read sequencing of males that produced extreme female biased sex ratios. Nucleotide diversity (pi) estimated from pool-seq WGS libraries is lower across the X for SR males. Compared to between replicate comparisons, differentiation (F_ST_) is only slightly elevated between populations with the same sex ratio type (between populations) but F_ST_ notably elevated chromosome-wide in pairwise comparisons between ST and SR pool-seq. Locations of named genes that were differentially expressed (DE) in testes between SR and ST RNAseq samples, or have differential coverage (DC) between SR and ST pools, or both (DCDE) are also shown. The shaded area indicates a 7MB region where five of the six inversions overlap, and includes reduced variation in all libraries, high differentiation, and a partial duplicate of *JASPer* that exhibits the strongest pattern of DC and DE among annotated genes.

Using long reads generated from *sex ratio* male siblings aligned to the reference genome, we identified six large X-chromosomal inversions that differ between SR males and the reference genome (**Figure 2**), which we further validated by examining short-read pool-seq data at break-point regions (**Table S3**). These inversions spanned the entire 97 MB chromosome except for a small region at the proximal end (0-2.04MB) and a region between 60.4 MB and 82.7MB. Many of the inversions overlap, particularly near the proximal end. Five of the six inversions are derived in the SR lineage and one (inversion 2) of six arose in the ST lineage when compared to the *Td2* draft genome (**Table S3**).Patterns of nucleotide diversity and differentiation appear to be influenced by proximity to the inversions. F_ST_ was elevated, and polymorphism reduced where there is a higher density of overlapping inversions. In particular, the region between 17MB and 20MB, where 5 of the 6 inversions overlap, has high F_ST_ between X^SR^ and X^ST^ and reduced diversity within all pools (**Figure 2**).

### Gene expression and copy number variation reveal drive candidates on the sex-ratio X

Using the pool-seq reads from X^SR^ and X^ST^ males above in combination with X^SR^ and X^ST^ RNAseq reads previously obtained from pools of mature testes (Reinhardt et al. 2014), we jointly evaluated differences in copy number and expression between X^SR^ and X^ST^. We identified 596 DE genes (37.6% had higher expression in X^SR^) and 120 genes with differential genomic coverage (DC genes, 62.5% with higher coverage on X^SR^). Among the 48 genes that were *both* DE and DC, there was a highly significant association in the direction of DE and differential genomic coverage (**Table S4**, FET p < 0.0001) with only 4 genes having higher X^ST^ expression and higher X^SR^ coverage. This result suggests differences in copy number often directly impact expression levels. As expected, most differences in gene expression (75.6%) and genomic coverage (89.2%) were confined to the X.

Seven annotated protein-coding genes (*JASPer*, *Pbp95*, *Tetraspanin 29Fb*, *Minichromosome maintenance 10*, *isopeptidase-T-3*, *Cyclin K*, and *santa-maria*) exhibited differential expression (DE) and differential genomic coverage (DC) between SR and ST males (**Table S4**). Strikingly, all seven were X-linked and had a higher level of both expression and coverage in *sex-ratio* males. The highest level of DE and DC was found for an X-linked paralog of *JASPer* (Jil-1 Anchoring and Stabilizing Protein), a gene which normally regulates the maintenance of euchromatin (Albig et al. 2019; Dou et al. 2020). Examination of genomic coverage near the *JASPer* region (X:18.18Mb) shows a 20-fold increase in coverage over a 1.5Kbp region containing a 1.1Kbp gene (**Figure 3A**). Long and short-read sequences from SR males showed supplemental alignment to nearby regions of the X at 18.19Mbp and 18.27Mbp. These two regions also contain a ~1.5 Kbp region with 20-fold increased raw short-read coverage. At 18.19Mbp the gene exhibits the same two-exon structure and length as the copy at 18.16Mbp but is positioned in the opposite orientation. At 18.27Mbp, a smaller (~600bp) single exon region is transcribed. Translations of the transcripts produced by these three partial *JASPer* genes contain only one of the two major functional domains of *D. melanogaster JASPer*, PWWP, which normally interacts with activating chromatin marks. There are also three full-length paralogs of *JASPer* on the C2 autosome and two additional X-linked copies that contain only the other functional domain, LEDGF, which interacts with *JASPer’s* functional partner, *JIL-1* (**Figure 3B**). Although eight other *D. melanogaster* proteins contain PWWP domains, a maximum likelihood phylogeny shows that the *T. dalmanni* JASPer PWWP domains are closer orthologs to the PWWP domain of *D. melanogaster JASPer*, than to other dipteran PWWP domains (**Figure 3C**), confirming these are partial *JASPer* paralogs. Neither of the LEDGF-only X-linked copies exhibit differential expression or coverage between SR and ST males (**Figure 3A**). The JASPer paralogs are all present in the reference genome at normal copy number but are amplified in copy number in X^SR^. Using only uniquely mapping SR and ST short reads, we estimated the increase in copy number of each JASPer copy on X^SR^. The fold-increase in genomic coverage of SR libraries across the amplified regions are ~58.9-fold (at 18.181:Mbp) and 3.4-fold (at 18.194:Mbp) and 6.5-fold (at 18.270:Mbp) compared to ST libraries (**Figure 3A**, “unique”). For the copy at 18.181:Mbp, we were also able to identify long reads which aligned uniquely to sequence matching both the left and right flanking regions and extended into the gene copy, and these contained five or six tandem copies of the PWWP-only *JASPer* paralog (**Figure S2**).

**Figure 3.**
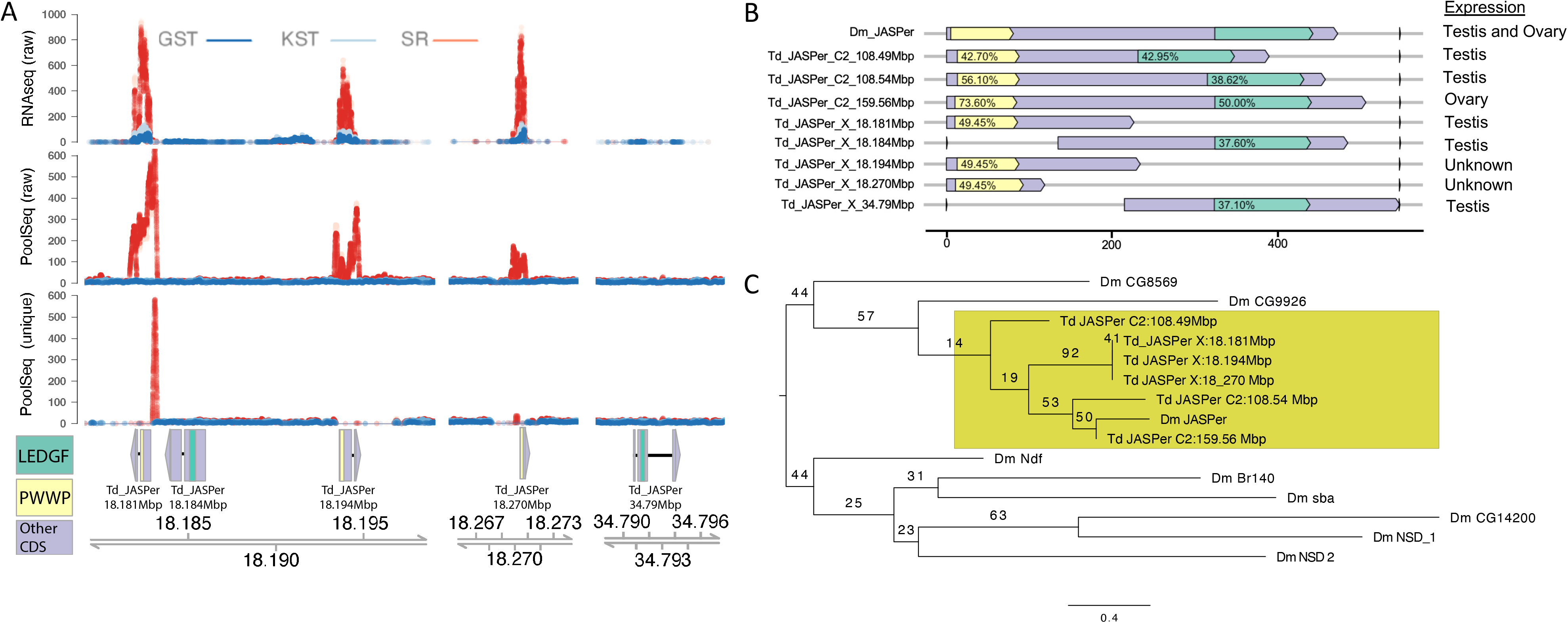
*JASPer*-PWWP has amplified in expression and covered on the sex ratio X (A) Examination of mapped reads in X-linked gene regions with homology to *JASPer* shows drastically higher expression (RNAseq) and coverage in pool-seq WGS libraries (pool-seq raw) from SR males, but not ST males from the same collections (GST and KST). Only those copies of *JASPer* including the PWWP domain showed increases in expression and coverage. Examination of uniquely mapping pool-seq reads (pool-seq Unique) demonstrates that the excess SR coverage is largely limited to the *JASPer* copy at 18.181 MB. (B) *T. dalmanni* contains eight transcripts with detectable homology to *D. melanogaster JASPer*. Copies on the second autosome (C2) contain both the two canonical domains (PWWP and LEDGF) whereas the five X-linked copies contain only one of the two domains. Amino acid percent identity compared to *D. melanogaster JASPer* is shown for each domain. (C) *T. dalmanni JASPer* copies are orthologous to PWWP from *D. melanogaster JASPer* as confirmed by a maximum likelihood phylogeny of all PWWP domains in *D. melanogaster* and all PWWP domains from *T. dalmanni JASPer* copies. The three X linked *JASPer* PWWP domains from ~18Mbp are identical in amino acid sequence.

### Transposable elements are enriched on the X and exhibit differential expression and copy number between X^SR^ and X^ST^

To investigate potential interactions between selfish genetic elements, we identified and annotated TEs across the *T. dalmanni* reference genome. Compared to *D. melanogaster*,repetitive sequences cover more of the *T. dalmanni* genome (~13.5% versus ~35.8%) (**Figure 4A**). About 10.0% of the genome is comprised of unclassified interspersed elements, while the rest includes 716 classified families from 38 superfamilies of Class I (DNA) elements (4.8% of genome) and Class II elements including LINE (10.7%) and LTR (5.9%) elements, but no SINE elements. Given the size of the *T. dalmanni* genome, this amounts to 14.4-fold as many insertions of classified TEs as in *D. melanogaster*, and 4.8-fold as many as the malaria mosquito *Anopheles gambiae*. LINE elements were significantly (χ^2^ = 3401, P<0.001) overrepresented in *T. dalmanni* compared to the other two species. The most abundant TE superfamilies in *T. dalmanni* include R1-LOA, Jockey, and RTE-BovB non-LTR (LINE) Class II elements (24,798, 15,719, and 11,712 copies, respectively), Gypsy LTR Class-II elements (14,484 copies) and TcMar-Mariner Class I (DNA) elements (13,210 copies).

**Figure 4.**
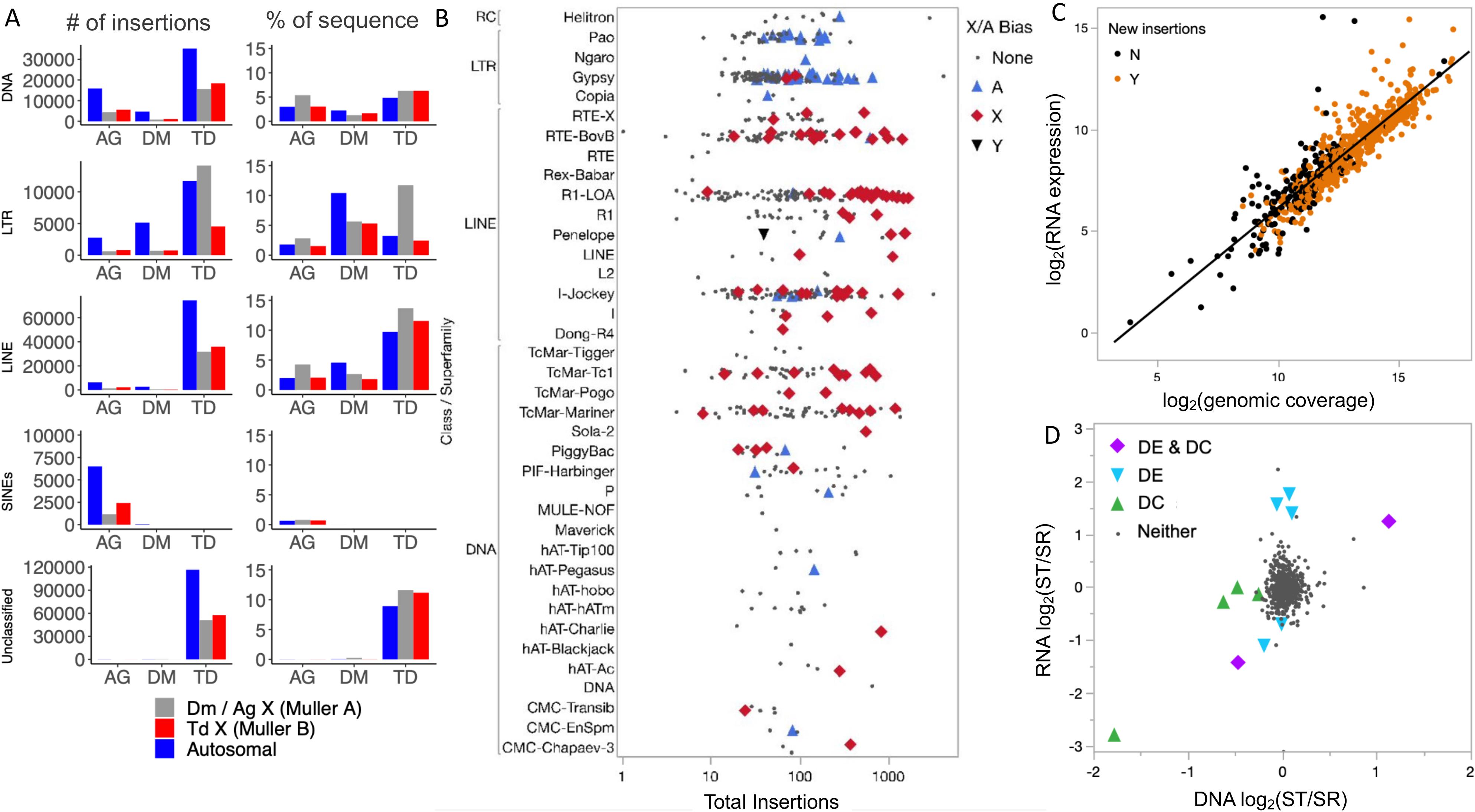
Transposable element (TE) colonization is ubiquitous across the genome in *Teleopsis dalmanni*. (A) The genome contains tens of thousands of insertions of all types of canonical elements except for SINEs, comprising 5-12% of the genome apiece, as well as many more interspersed elements that could not be classified. (B) There was a bias towards preferential insertion on the X chromosome for more superfamilies, especially Class-I non-LTR (LINE) and Class-II (DNA) elements. Multiple LTR superfamilies are autosome-biased, driven by a large number of insertions of Gypsy and Pao family elements on the distal arm of the first autosome (Muller element A). One Penelope element is enriched in male sequencing data compared to female resequencing data, indicating Y-enrichment. (C) TE families that are expressed at a higher level (as assessed by RNAseq) and present at a higher copy number (as assessed by quantification of PoolSeq WGS libraries) are more likely to have produced new, polymorphic insertions than rarer, less highly expressed elements. (D) Expression and/or copy number of eleven TE families is impacted by sex ratio status, with 5 of 6 elements with X^SR^ / X^ST^ differential coverage (DC) having higher copy number in the SR Pools. X^SR^ / X^ST^ differential expression (DE) was split between higher expression in SR and ST Pools.

Overall, transposable elements are more abundant on the *T. dalmanni* X than on the two autosomes (293.9 TEs/Mb on the X *vs*. 204.5 TEs/MB on the two autosomes, χ^2^=364.46, P<0.0001). TE distribution also varies by element type (**Figure 4A**), with more X-linked DNA elements (26.6%, χ^2^=364, *P*<0.0001) and LINE elements (25.3%, χ^2^=372, *P*<0.0001) but fewer X-linked LTR elements than expected (14.9%, χ^2^=532, *P*<0.0001) given the X comprises 22.2% of the genome. Transposable elements are found more often than expected on *T. dalmanni* Muller A, and less often than expected in the comparison species where Muller A is an X chromosome (χ^2^= 656, P<0.0001). This appears to be driven by an excess of LTR elements on Muller element A – 42% of all LTR elements are found on this single chromosome arm in *T. dalmanni*. We found that 28 of 184 Class I (DNA) families have more copies (**Figure 4B**) than expected on the X chromosome (χ^2^= 11.59, P < 0.0001) including 19 different TcMar families, while 61 of 337 LINE families have more copies than expected on the X (χ^2^= 11.59, P < 0.0001). In contrast only 2 of 184 LTR element families are enriched on the X (χ^2^= 65.10, P < 0.0001), while 50 of 184 LTR families are more commonly autosomal. Most of these autosome-enriched LTRs are Gypsy and Pao elements (**Figure 4B**). One Penelope element was present on male reads 8-fold more than on female reads, consistent with Y-amplification, and several other elements also were male-biased. Among the top 10 most male-biased families were two other Penelope elements, five Gypsy LTR elements, and two Pao LTR elements (**Table S5**).

We also found evidence for robust expression of many TE families in the *T. dalmanni* genome (**Figure 4B**). The activity of each TE family was assessed using SR and ST testis RNAseq (Reinhardt et al. 2014), and by identifying novel insertions present in one or more of the genomic resequencing pools. Across TE families, normalized genomic coverage correlated strongly with normalized expression in standard males (**Figure 4C**). Unsurprisingly, TE families that were highly expressed and abundant in the genome were more likely to produce new insertions than those with low expression and genomic abundance (**Figure 4C**). We assessed SR-ST differential expression and SR-ST differential coverage of the 716 classified TE families and did not find a correlation between SR-ST differential expression (DE) and SR-ST differential coverage (DC) (**Figure 4D**). Seven TE families were differentially expressed (DE) between SR and ST testis (3 up in SR and 4 up in ST) and six exhibited differential genomic coverage with five present at a significantly higher copy number (DC) in SR male genomic DNA (**Figure 4D**, **Table S5**). Compared to a 1:1 expectation, we find no difference between the number of families that are either DE (χ^2^=1.47, P =0.41) or DC (χ^2^=2.667, P =0.26) in SR versus ST testis.

### Genome defense genes are diverging under positive selection, including between X^ST^ and X^SR^

Given the high levels of TE expression and evolution we observed, we investigated the evolution of genome defense genes in the *T. dalmanni* genome. To predict *dN/dS* between *T. dalmanni s.s*.and *T. dalmanni sp 2* we fit a series of linear models that included one or more of the following variables: involvement of a gene in the piwi-interacting RNA (piRNA) genome defense pathway, tissue expression pattern, and X-linkage. The best-fitting model (F-stat=132.4, df=4/5592,*p* < 2.2 e-16, R^2^ = 0.086) revealed a significant interaction between testis expression and X-linkage (**Figure 5A**), and accelerated evolution of piRNA pathway genes (**Figure 5B, Table S6-S7**). Rates of evolution are fastest among X-linked genes expressed in testes that are involved in the piRNA pathway. *Aubergine* (*aub*), a piwi gene family member that complexes with antisense RNA during TE silencing, is the fastest evolving with *dN/dS* = 1.234. This fast-evolving *Aubergine* is a testis-specific paralog of the gene; an ovary-specific, slower evolving (*dN/dS* = 0.104) *Aub* paralog is also present. McDonald-Kreitmann tests performed on a set of core piRNA related genes further identified *spn-E* as putatively evolving under positive selection in *Teleopsis* (**Table S8**).

**Figure 5.**
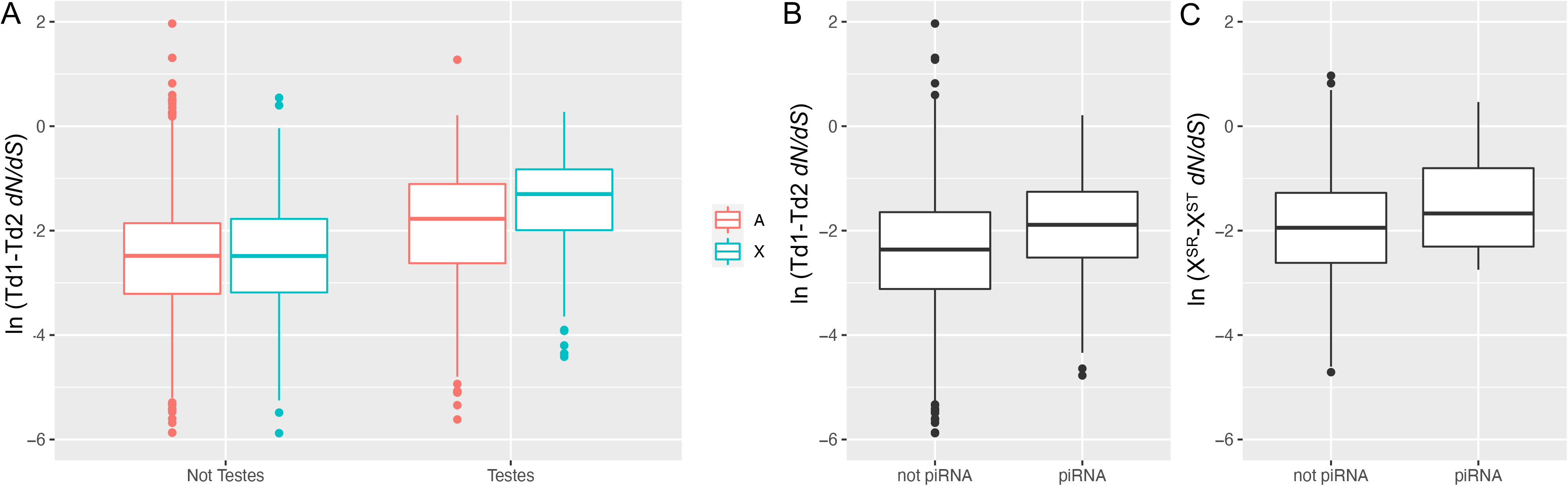
Rapid evolution of piRNA related genes in *Teleopsis dalmanni. dN/dS* between *T. dalmanni* sp1 and *T. dalmanni* sp 2 (Td2) was assessed for 9,525 protein-coding genes and *dN/dS* between the X^SR^ and X^ST^ chromosomes was assessed for 2,642 genes. Significance of comparisons was assessed using linear models on log-transformed *dN/dS* values and AICc model selection (Table S6-S9). The expression pattern of genes was annotated based on tissue-specific RNAseq in *T. dalmanni* (Baker 2016). Linkage was determined as either X-linked (X) or Autosomal (C1 or C2 chromosome) by alignment to the genome assembly with GMAP. The status of a gene as “piRNA related” was determined based on prior work in *Drosophila* (Palmer 2018, Tables S1 and S2). (A) piRNA related genes evolved faster between *T. dalmanni s.s*. and *Td2* (B) Genes evolved significantly faster when they were testis expressed and X-linked (there was a significant interaction between linkage and expression pattern). (C) X-linked piRNA related genes also evolved faster between sex-ratio and standard males.

Sixteen of 80 RNA interference orthologs (14 piRNA related, 1 siRNA related, and 1 miRNA related) are X-linked in *T. dalmanni* and therefore we measured divergence and molecular evolution between X^SR^ and X^ST^ alleles of the genes (**Figure 5C, Table 1**). We found seven RNAi genes (**Table 1**) that were differentially expressed (DE) between X^SR^ and X^ST^ and the direction of change varied by mode of RNA interference. Two genes involved in siRNA function (*vig* and *dcr-2*) were upregulated in ST testes, whereas five piRNA related genes had higher expression in SR testes. Multiple piRNA-related genes have X-linked paralogs of genes that are single copy in *D. melanogaster*. For example, *Maelstrom* (*mael*) is present in five copies within the *T. dalmanni* genome, four of which arose within the genus (**Figure S3A**). The X-linked and testis-specific *maelstrom* paralog is evolving under positive selection between X^SR^ and X^ST^ (*dN/dS* = 1.58) and has six nonsynonymous X^SR^-X^ST^ differences, including several within its self-named functional domain (**Figure S3B**).

**Table 1.**
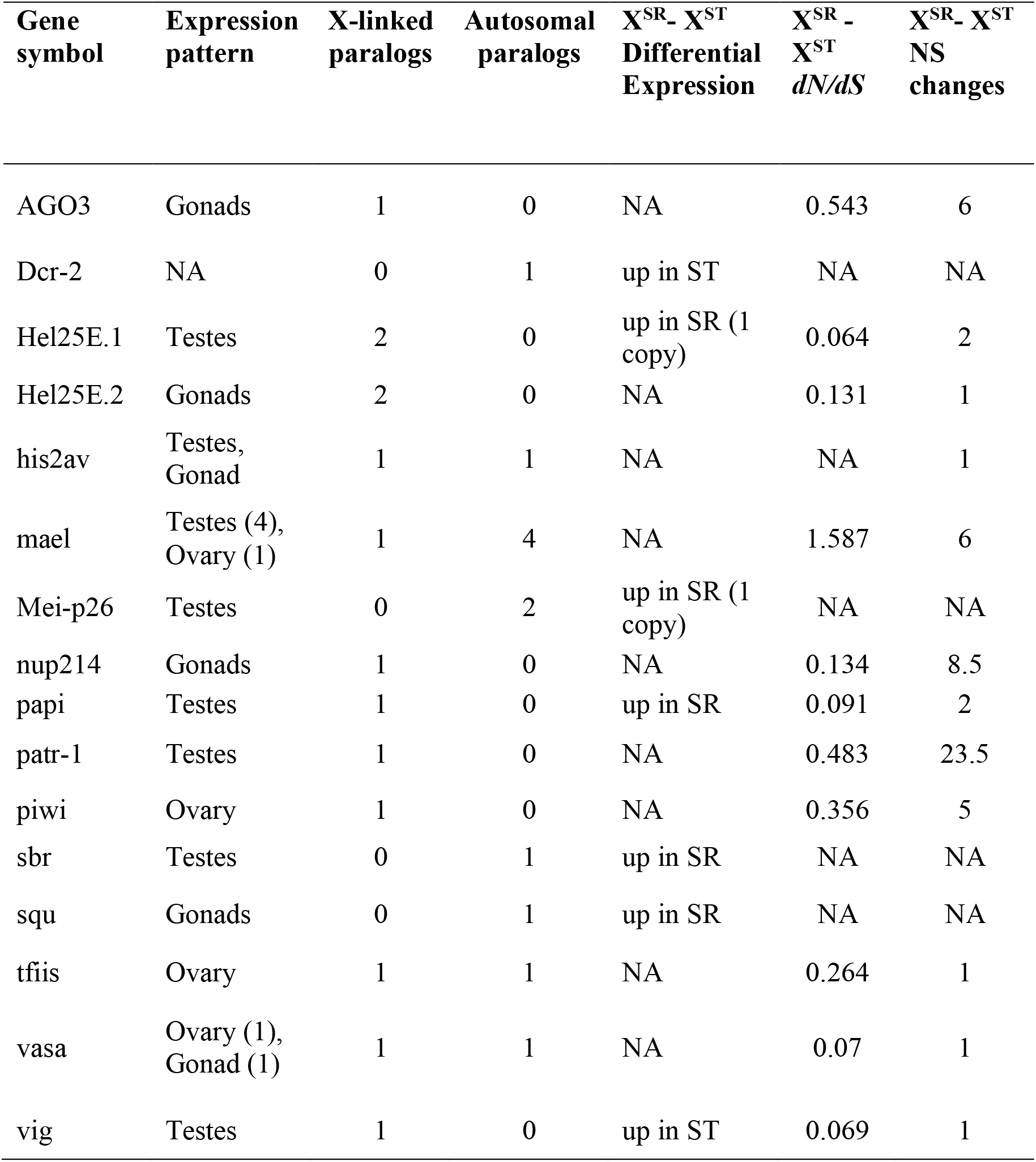
Impacts of X^SR^ and X^ST^ divergence on X-linked RNAi pathway genes.

## Discussion

Sex-linked meiotic drivers are selfish genetic elements that can lead to dramatic genomic changes as their interests conflict with those of other genes (Burt and Trivers 2006). Here, we analyzed the impacts on a stalk-eyed fly (*Teleopsis dalmanni*) genome of a long-term association with such an element. Prior work suggested widespread differentiation between X^SR^ and X^ST^ chromosomes (Reinhardt et al. 2014) and identified at least one inversion distinguishing the two (Paczolt et al. 2017). By assembling the genome into three chromosome-length scaffolds we find dramatic differentiation between X^SR^ and X^ST^ that extends across the entire X chromosome, a sign that genetic recombination between the two types of X chromosomes has been severely limited for a long time (**Figure 1, Figure 2**). By combining pool-seq short-read and long-read resequencing data we located the break points of six overlapping inversions that span most of the X (**Figure 2, Table S3**) and have permitted sequence divergence to persist between the inversion haplotypes. We identified several genes that have diverged dramatically in copy number and expression between X^SR^ and X^ST^ including a paralog of *JASPer*, a gene known to be involved in regulation of chromatin and TE expression, and the key TE control gene *maelstrom*. Transposable elements have also proliferated differentially on X^SR^ and X^ST^, suggesting these effects may not be independent.

Work in *Drosophila* over the last few decades has implicated genome defense and / or the regulation of heterochromatin as common denominators underlying meiotic drive (Courret et al. 2019). Most meiotic drive systems with a known molecular mechanism involve regions of repetitive DNA either in the target of drive (Wu et al. 1988; Tao et al. 2007b,a; Helleu et al. 2016) or in both the driver and its target (Hurst 1996; Cocquet et al. 2012) that may be targeted by genome defense. For example, in the *Segregation Distorter (SD) – responder* system in *D. melanogaster*,small RNA matching the 240 bp *responder* (Wu et al. 1988; Houtchens and Lyttle 2003; Larracuente 2014) repeats has been shown to be associated with piRNA pathway genes, *Aubergine* and *Argonaute3* (Nagao et al. 2010), and when *Aubergine* was knocked out in an *SD* background, the strength of distortion by the driver was enhanced (Gell and Reenan 2013). The first piRNA locus to be characterized, *Suppressor of Stellate* (Aravin et al. 2007) likely first originated as a suppressor of drive (Hurst 1996), and now acts to silence TEs. And more recently, *Ago-2* mediated hpRNA/endogenous siRNAi pathway (Chung et al. 2008; Czech et al. 2008; Ghildiyal et al. 2008; Kawamura et al. 2008) has been implicated in the Winters and Durham sex ratio meiotic drive systems in *D. simulans* (Lin et al. 2018; Muirhead and Presgraves 2021; Vedanayagam et al. 2021).

We find the strongest candidate for drive in this species to be the *JIL-1 Anchoring and Stabilizing Protein (JASPer*, also known as *dP75). JASPer* has multiple new paralogs in the genus *Teleopsis*, and furthermore, partial X-linked copies have amplified in expression and copy number on X^SR^ relative to X^ST^ (**Figure 3, Table S4**). In *D. melanogaster, JASPer* positively regulates the transition of heterochromatin to euchromatin with its partner JIL-1 (Albig et al. 2019) and is essential during oogenesis (Dou et al. 2020). A partial paralog of *JASPer* appears to have formed tandem arrays on X^SR^, with only part of the gene – the chromatin binding PWWP domain - being duplicated and upregulated in expression in X^SR^ (**Figure 3**). This amplified partial paralog (*Td-JASPer:X:18.181MB*) is found in a region overlapped by 5 of the 6 inversions we identified as distinguishing X^SR^ and X^ST^ (**Figure 2**), and is also testis-specific, whereas the full-length paralog closest in sequence to the *D. melanogaster* ortholog (at *Td-JASPer:C2:156MB*) is primarily expressed in the ovary (Figure 3B). PWWP domains, like those found in *JASPer*, primarily bind to H3K36me3 chromatin marks (Albig et al. 2019; Dou et al. 2020), which associate with regions of active gene expression. Although we do not know the target of any *JASPer* paralog in *T. dalmanni*, it would seem likely this function is conserved given it occurs across Metazoa. Plausibly, the PWWP-only *JASPer* duplicates expressed from X^SR^ might bind to their targets but are unable to recruit *JIL-1* and therefore fail to properly activate target genes, similar to a dominant negative allele. If the chromatin targets of *PWWP-JASPer* are primarily Y-linked, development of Y-bearing, but not X^SR^-bearing, sperm could be interrupted. *JIL-1/JASPer* binding is also implicated in positive regulation of expression of *Gypsy5* retroelements, which are found in arrays in *Drosophila* telomeres (Albig et al. 2019), providing a plausible connection between this drive candidate and disruption in expression of certain TEs. We found that Gypsy elements are generally less common on the X chromosome, and several Gypsy families show a male bias in genomic coverage suggesting amplification on the Y (**Table S5**). PWWP-*JASPer* duplicates could even be targeting these Y-specific elements to cause meiotic drive.

We also found multiple families of TEs (**Figure 4B**) and several genes involved in genome defense (**Table 1**) that are diverging in expression and copy number on the X^SR^ chromosome. X-linked piRNA pathway genes including *piwi*, *Argonaute3*, and an X-linked copy of *maelstrom* have accumulated nonsynonymous differences compared to the heterokaryotypic allele on X^ST^ (**Table 1**), with *maelstrom* showing evidence of positive selection between X^SR^ and X^ST^ (**Figure S3**). Meanwhile, complete genome assembly allowed a comprehensive transposable element annotation revealing over 261,000 TE copies comprising ~25% of the genome. X^SR^ associated differences in TE expression and copy number may be caused by genetic drift between the two chromosome types – X^SR^ has reduced recombination rates and polymorphism, subjecting the chromosome to effects of Muller’s Ratchet (Muller 1964). Prior work has shown that some piRNA pathway proteins, like immune genes, evolve rapidly (Obbard et al. 2009; Kolaczkowski et al. 2011; Lee and Langley 2012; Simkin et al. 2013; Palmer et al. 2018), although the mechanism underlying this rapid evolution is unclear (reviewed in (Blumenstiel et al. 2016), but see (Parhad and Theurkauf 2019). We, too, find that piRNA genes are generally more rapidly evolving than other genes (**Figure 5, Table S6-S7**) and some of these genes (*spn-E* and *aubergine*) show signs of evolving under positive selection within *T. dalmanni* (**Table S8**). A missing piece here is the expression of the piRNA themselves. Given that piRNA clusters are formed via new mutations (Zhang et al. 2020), if present on the X, they should accumulate different TE insertions over time due to the lack of recombination between X^ST^ and X^SR^ and so target different mobile elements. *Maelstrom’s* role in TE silencing involves promoting the spread of heterochromatic states to nearby genes (Sienski et al. 2012) so its adaptive X^SR^ and X^ST^ divergence could either be a symptom or contributor to meiotic drive, perhaps in concert with JASPer.

Overall, these results extend the work of others on Drosophila (reviewed in Courret et al. 2019) to another charismatic Dipteran – *Teleopsis* stalk-eyed flies – and suggest that associations among meiotic drive, regulation of heterochromatin, and the regulation of transposable elements by RNAi may be common in Diptera (Hurst 1996; Nagao et al. 2010; Gell and Reenan 2013; Larracuente 2014; Lin et al. 2018; Muirhead and Presgraves 2021; Vedanayagam et al. 2021), even as the details of molecular mechanisms vary.

## Material and Methods

### Genome assembly of *Teleopsis dalmanni s.s*

A draft genome assembly for *Teleopsis dalmanni*, NLCU01 was created with a combination of Roche 454, Illumina and Pacific Biosciences sequence data (**Table S1**) using MaSuRCA (Zimin et al. 2013) and is available on Ensembl metazoa, GCA002237135v2 (Kersey et al. 2018). This assembly contains ~25k scaffolds with an N50 of ~75k. All DNA sequences were obtained from an inbred population (line “2A”) of *Teleopsis dalmanni*. This population is derived from flies that were first collected near the Gombak River in peninsular Malaysia (3 12’N, 101 42’E) in 1989 and then maintained as a control line for an artificial selection study on relative eyespan (Wilkinson 1993; Wolfenbarger & Wilkinson 2001). After 50 generations of selection, full-sib mating was conducted for seven generations to establish the line, which has subsequently been maintained without additional inbreeding. This population has been used in several prior studies (Christianson et al. 2005; Wilkinson et al. 2014) and does not carry any drive-associated genetic markers. Contaminating bacterial scaffolds were identified and removed prior to submission using a modification of the (Wheeler et al. 2013) DNA-based homology pipeline (Poynton et al. 2018). Male and female genomic short-read resequencing data was aligned to each scaffold using nextgenmap (Sedlazeck et al. 2013) with default parameters, and relative coverage of male and female reads was used to identify X-linked scaffolds (cf. Vicoso and Bachtrog 2015), with the expectation that the normalized ratio of female to male reads should be approximately 1 to 1 for autosomal and 2 to 1 for X-linked scaffolds. The initial assembly was filtered for haplotigs and other redundant sequences using Purge Haplotigs (Roach et al. 2018).

A chromosome-level assembly (NLCU02) was then created by incorporating chromatin conformation information and validated with a high-density linkage map. Chromatin conformation capture data was generated using a Phase Genomics (Seattle, WA) Proximo Hi-C Plant Kit, which is a commercially available version of the Hi-C protocol (Lieberman-Aiden et al. 2009). Following the manufacturer’s instructions, intact cells from unsexed pupae from the 2A inbred line were crosslinked using a formaldehyde solution, digested using the Sau3AI restriction enzyme, and proximity-ligated with biotinylated nucleotides to create chimeric molecules composed of fragments from different regions of the genome that were physically proximal in vivo, but not necessarily genomically proximal. Continuing with the manufacturer’s protocol, molecules were pulled down with streptavidin beads and processed into an Illumina-compatible sequencing library. Sequencing was performed on an Illumina HiSeq 4000, generating a total of 202,608,856 100 bp read pairs. Reads were aligned to the draft assembly (NLCU01.30_45_breaks.fasta) following the manufacturer’s recommendations. Briefly, reads were aligned using BWA-MEM (Li and Durbin 2009) with the −5SP and -t 8 options specified, and all other options default. SAMBLASTER (Faust and Hall 2014) was used to flag PCR duplicates, which were later excluded from analysis. Alignments were then filtered with samtools (Li et al. 2009) using the -F 2304 filtering flag to remove non-primary and secondary alignments. Putative misjoined contigs were broken using Juicebox (Rao et al. 2014; Durand et al. 2016) based on the Hi-C alignments. A total of 113 breaks in 105 contigs were introduced, and the same alignment procedure was repeated from the beginning on the resulting corrected assembly. The Phase Genomics’ Proximo Hi-C genome scaffolding platform was used to create chromosome-scale scaffolds from the corrected assembly as described (Bickhart et al. 2017). As in the LACHESIS method (Burton et al. 2013), this process computes a contact frequency matrix from the aligned Hi-C read pairs, normalized by the number of Sau3AI restriction sites (GATC) on each contig, and constructs scaffolds in such a way as to optimize expected contact frequency and other statistical patterns in Hi-C data. In addition to Hi-C data, chromosomal linkage information (see below) was used as input to the scaffolding process. Linkage groups from a linkage map were used to constrain chromosome assignment during the clustering phase of Proximo by discarding any suggested clustering steps that would incorporate contigs from different linkage groups onto the same chromosome, but linkage map data were not used during subsequent ordering and orientation analyses in Proximo. Approximately 528,000 separate Proximo runs were performed to optimize the number of scaffolds and scaffold construction to make the scaffolds as concordant with the observed Hi-C data as possible. This process resulted in a set of three chromosome-scale scaffolds containing a total of 438.2MB, comprising 95.7% of the filtered MaSuRCa assembly. Finally, Juicebox was again used to correct remaining scaffolding errors.

Linkage groups used to validate the Hi-C assembly were created by mating a female hybrid offspring obtained from a cross between a male from the 2A inbred strain and a female from a noninbred population of *T. dalmanni* collected near Bukit Lawang, Sumatra (3 35’N, 98 6’E) to a male from the 2A strain. This backcross produced 249 (131 female and 118 male) individuals which were individually genotyped using multiplex shotgun sequencing (Andolfatto et al. 2011) and multiple STR loci (Wilkinson et al. 2014). Genotypes were determined as either heterozygous or homozygous for each scaffold by combining all loci present on a scaffold into a single “super locus”. Reads were aligned using bwa (Li and Durbin 2009) and genotypes were assessed as either homozygous or heterozygous using samtools v.1.9 (Li et al. 2009) (mpileup -v). Because this was a backcross, for autosomal loci those individuals with the backcross allele (pure “2A”) should be homozygous at informative markers whereas individuals with the non-backcross allele should be heterozygous, with an expectation of a 1 to 1 ratio of these genotypes. This results in an overall 3 to 1 ratio of the backcross to the non-backcross allele for autosomal markers and X linked markers in females, and an overall 1 to 1 ratio for X linked markers in hemizygous males. Markers were retained as potentially informative if at least one individual was found to carry the non-backcross (Wilkinson et al. 2014) allele, which we defined as the less common allele across all female individuals. Markers were removed if they violated expected allele ratios for a backcross using a binomial test against the expectations described above, or if more than 20% of female individuals were found to carry **only** the non-backcross allele. Finally, within each individual all markers from a given scaffold were pooled to give an overall number of reads supporting each genotype and requiring a minimum coverage of 5 reads per marker. Individuals were assigned in the final matrices as “a” (for 2A/backcross genotypes) or “b” (for Bukit Lawang/foreign genotypes).

Separate genotype matrices were then created for the X-linked scaffolds (as determined by male and female coverage (Vicoso and Bachtrog 2015) and autosomal scaffolds and rank-ordered by the number of individuals genotyped. We then used JoinMap (Stam 1993) v4.1 to assign the top 1000 autosomal scaffolds into one of two linkage groups (chromosomes). Only scaffolds with a LOD score > 5 were assigned to a chromosome. We used a similar process for the top 250 X-linked scaffolds but used a LOD score > 10 to assign scaffolds to the chromosome. These linkage groups were used to constrain the Hi-C genome assembly as noted above. Then, independent from the Hi-C scaffolding process, we ordered scaffolds within each linkage group by regression mapping using a Haldane mapping function (Haldane’s Mapping Function 2008). We used regression mapping, rather than maximum likelihood, because it is less sensitive to missing genotype data (Van Ooijen 2006) as is typical for MSG studies. We removed markers from the final map if there was evidence for significant (after Bonferroni correction) lack of fit to their nearest neighbors. The resulting linkage map included 762 scaffolds spanning 147.1 Mbp. Collinearity between the Hi-C and linkage maps was assessed by comparing the relative position of scaffolds which were found in both maps. Chromosomal synteny of the assembly scaffolds with *Drosophila melanogaster* chromosome arms was assessed by alignment of a set of 7,634 previously annotated 1-to-1 *Drosophila* orthologs (Baker et al. 2016) to the genome assembly using GMAP (Wu and Watanabe 2005) (--npaths=1 --format=gff3_gene --min-identity=0.9). Assemblies were assessed for completeness using BUSCO (Seppey et al. 2019) against 3,285 conserved dipteran genes.

### Draft Genome assembly of *T. dalmanni* species 2

We previously described a cryptic species of stalk-eyed fly (Paczolt et al. 2017), which we refer to as *T. dalmanni sp 2* (Td2) and which corresponds to prior collections of *T. dalmanni* from several sites in peninsular Malaysia, such as Cameron Highlands (Christianson et al. 2005; Swallow et al. 2005). To produce a draft genome of this species, we extracted HMW DNA from a single male from a laboratory population of Td2 using the Gentra Puregene tissue kit (Qiagen cat 158667). 1 ug of DNA was sent to the New York Genome Center (NYGC) where it was prepped with the Chromium Genome linked read kit (10X Genomics) and sequenced on a half lane of an Illumina HiSeqX machine, producing a total of 416 million reads. These reads were assembled at NYGC using Supernova (v2.0.1) (Weisenfeld et al. 2017). The resultant draft genome contained 10,290 scaffolds greater than 10 kb with a N50 of 45.2 kb and total genome size of 355 MB. While incomplete, this genome was sufficient to determine inversion history, and polarize molecular divergence.

### Short-read resequencing of sex ratio (SR) and standard (ST) males

To identify sequence and structural variations specific to X^SR^, we sequenced DNA from replicate pools of sex ratio (SR) males (males with female-biased offspring sex ratios), or standard (ST) males either collected in the field from two different sites in peninsular Malaysia or representing the first three generations of sons descended from field-collected females. One SR and two ST sample pools were created from the DNA of males from each of two collection sites (Gombak and Kanching) that were previously phenotyped for offspring sex ratio (Paczolt et al. 2017). When an excess of individuals was available from a collection and *sex ratio* category, genotype data from nine X-linked STR loci from the same analysis was used to avoid over-sampling closely related individuals (e.g. male full-siblings from the same brood). Consequently, haplotype diversity within pools was not significantly different to haplotype diversity among all candidate males for that pool (following (Christianson et al. 2011), Dunnett’s t-test, p>0.05 for all comparisons, **Table S9**). DNA was extracted using the DNeasy Blood and Tissue Kit (Qiagen) and quantified using PicoGreen Quant-IT dsDNA quantification kit (Thermofisher Q33130). Pools were then assembled using an equimolar amount of DNA from each sample. Sample size for each pool ranged from 15-18 individuals (**Table S9**). Six barcoded libraries were prepared and multiplexed on two lanes of a HiSeq1500 set to RapidRun mode to generate 150 bp paired-end sequences. Bam-formatted alignments of these libraries to the genome were produced using nextgenmap (Sedlazeck et al. 2013) with default parameters and used in subsequent analyses. Pairwise genetic diversity for each pool and FST between each pairwise combination of pool-seq data was calculated from the pooled resequencing alignments in 5kb non-sliding windows using popoolation2 (Kofler et al. 2011). After pool-seq had been completed, it was determined that four pooled ST individuals were actually *Td2* males. Genetic markers distinguishing these species had not been identified until after pooling, see **Table S9**, (Paczolt et al. 2017), and previous work (Christianson et al. 2005; Swallow et al. 2005) had suggested that *Td2* would not be present in the collection sites we visited. Both SR pools and one ST pool (Kanching ST2) were entirely composed of *T. dalmanni* s.s. (**Table S9**) so all analyses where excess polymorphism (potentially caused by species divergence from the *Td2* individuals) within a pool could impact the results of analysis were repeated using only these three samples, and results of the reduced analysis were found to be qualitatively similar to the full analysis (**Table S10**).

### Long-read resequencing of a SR haplotype

To identify inversion breakpoints between X^SR^ and X^ST^, we used long-read sequencing (Pacific Biosciences). A pool of full-sib males bearing a single identical-by-descent (IBD) X^SR^ haplotype was created by mating a SR/SR female to a male from the 2A strain and then backcrossing the female progeny to males from the same strain. We then genotyped 107 sons from this backcross at three X-linked STR loci (ms125, ms395, and CRC) to distinguish X^SR^ and X^ST^ sons. DNA from 46 X^SR^ sons was then extracted using the Gentra PureGene Tissue Kit (Qiagen) and pooled, followed by a phenol-chloroform extraction and ethanol precipitation. A PacBio long insert (15 Kb) library was then prepared and run on three PacBio Sequel SMRT cells. These runs yielded a total of 15.1 Gb of sequence, with mean read length of 4.9 Kb and maximum read length of 92.8 Kb. Raw long reads were aligned to the genome using ngmlr with default parameters and structural variants were called using sniffles (Sedlazeck et al. 2018), requiring at least 2 reads to support each variant call. The resulting output was filtered to find inversions that were fixed within the SR PacBio long reads relative to the reference genome. Each putative inversion was then validated as a fixed SR specific inversion by comparison to the read-pair orientation in the SR and ST pool-seq data at the breakpoint using IGV (Thorvaldsdottir et al. 2013). An inversion was considered validated if reads from the SR pool-seq samples but none of the ST pool-seq samples agreed with the sniffles call at that position (**Table S3**). Finally, to polarize the direction of the inversion mutation, an alignment of the *T. dalmanni sp 2* genome assembly was performed using blat (Kent 2002) and scaffold alignments near the breakpoints were examined to determine if they 1) support the standard arrangement (span the breakpoint), 2) support the SR arrangement (scaffold breaks and aligns to other end of breakpoint), 3) support another arrangement (scaffold present near breakpoint), or 4) are uninformative (no scaffolds map near the breakpoint) (**Table S3**).

### Transposable element annotation

Transposable elements were annotated in the *T. dalmanni ss* assembly using RepeatModeler v. 1.0.4 (Smit and Hubley 2008) with default parameters and the NLCU02 assembly as input. Resulting consensus fasta formatted TE sequences were input into RepeatMasker (Smit et al. 2013) with the assembly as the reference, producing a repeat-masked reference genome and repeat annotations. The tool “One code to find them all” (Bailly-Bechet et al. 2014) was used with the RepeatMasker output (.out) to count the numbers and locations of each type of insertion in the *T. dalmanni* genome. These were compared to the RepeatMasker annotations for two other dipterans (*Drosophila melanogaster* dm6 RepeatMasker open-4.0.6 and *Anopheles gambiae* anoGam1 RepeatMasker open-4.0.5) analyzed using the same procedure. Further classification of element superfamilies followed prior universal classification schemes (Wicker et al. 2007; Makałowski et al. 2019). Sex (or autosomal) chromosome bias in the distribution of elements of each annotated family was determined by comparing the observed number of intact elements of each element type on the X to the number expected assuming the X comprises 22.5% of the genome using a Chi-squared goodness-of-fit test. To correct for multiple testing, we applied a Benjamini-Hochberg (BH) 5% False Discovery Rate (FDR).

Novel, polymorphic insertions of TEs were called in each of the six Illumina resequencing pools using PopoolationTE2 (Kofler et al. 2016) running the “separate” analysis mode on the repeat masked assembly and TE consensus sequences. Sites were subsampled to 20x coverage and were discarded if coverage was less than 20x in any sample. To compare the rate of insertions of TEs between the samples, insertions were inferred to be orthologous if they were an insertion of the same element within 500 bp in multiple pools.

The expression of RepeatModeler TE families in SR and ST male testes was assessed using TEtools (Lerat et al. 2016), using the default settings and including alignment with bowtie2 (Langmead and Salzberg 2012) using the RepeatModeler TE library and RNAseq reads from two pools of SR male testis and two pools of ST male testis (BioProject PRJNA240197). Unannotated repetitive elements (“Unknown” interspersed and simple repeats) were removed after normalization but prior to differential expression analysis with DESeq2 (Love et al. 2014). Differential TE expression between the SR and ST pools was assessed using the negative binomial Wald test on the DESeq-normalized counts for each TE family. TEtools was also used to estimate TE family copy number within the SR and ST genomic resequencing pools and in male and female genomic libraries (SRS2309195-SRS2309198). We identified potentially Y-inserted elements by qualitative comparison of male and female DESeq2 normalized genomic read counts (lack of replication prevented statistical analysis).

### Annotation of gene duplication and expression, and differential coverage

A set of protein-coding genes annotated from a transcriptome assembly (BioProject PRJNA240197) was aligned to the three largest scaffolds using GMAP allowing for up to 10 gene alignments (“paths”) per gene (--npaths=10 --format=gff3_gene). Annotations were removed as potential TEs misannotated as genes if they had >50% alignment overlap with any TE annotation from RepeatMasker. BEDTools (Quinlan and Hall 2010) (intersect -wao) was used to determine the number of bases of overlap for each exon, then the proportion of overlapping bases was calculated across the entire length of each gene alignment (“path” in GMAP terminology). RNAseq data from SR and ST male testis (Reinhardt et al. 2014) was aligned to the genome using HISAT2 (Kim et al. 2019) v2.0.1-beta (--dta -X 800). As previously described genomic pool-seq data were aligned to the genome using nextgenmap (Sedlazeck et al. 2013). Genomic expression and coverage in each library was estimated using Featurecounts (Liao et al. 2014). Differential coverage and differential expression were each assessed from the Featurecounts read count matrix on a by-feature basis using DESeq2 (Love et al. 2014) with the default Wald test on the negative binomial distribution.

### Molecular evolutionary analyses

Divergence was assessed in comparison to the *Teleopsis dalmanni sp* 2 (Paczolt et al. 2017) genome described above. The *Td2* draft genome scaffolds were aligned to the three largest (chromosomal) scaffolds in the Hi-C assembly for *T. dalmanni s.s*. using GMAP (Wu and Watanabe 2005) (--nosplicing --format=samse). A *Td2* consensus was called from the GMAP alignment of *Td2* scaffolds to the *Teleopsis dalmanni* genome by sorting and indexing with samtools (Li et al. 2009) v. 1.3.1, then calling the consensus with Bcftools (Li 2011) v 1.9 (bcftools call --ploidy 1 -mA). Regions which did not have an aligned scaffold or align as gaps show up as stretches of “N’s” in the consensus when these parameters are used.

For molecular evolutionary comparisons, the bam-formatted pool-seq library alignments were used with bcftools (bcftools call --ploidy 1 -c; vcfutils.pl vcf2fq), to create a majority-rule X^SR^ consensus sequence for the large X-chromosomal scaffold (PGA_scaffold1) from a bam file combining both X^SR^ pools into a single bam file. In addition, to have a comparable (similar sequencing and allelic coverage) X^ST^ consensus, an X^ST^ alignment (bam) file was produced using 1 replicate from each collection site (Gombak ST1 and Kanching ST2) and consensus called as above. The best alignment of the coding regions of genes previously (Baker et al. 2016) assembled and annotated using data from a multi-tissue RNAseq experiment (BioProject PRJNA240197) were localized to the X-chromosomal scaffold via alignment with GMAP (--npaths=1 -- format=gff3_gene --min-identity=0.9). Gene sequences were extracted from the X^SR^ and X^ST^ consensus X chromosomes described above using the GffRead utility (Pertea and Pertea 2020) and the mRNA gff annotations from GMAP. Some genes contained in-frame stop codons in one or more libraries - these genes were trimmed to the longest open reading frame present in all pools and if what remained was longer than 50 amino acids, were retained. Genes were also excluded if they contained only ambiguity sequence (“N”) in one or more of the consensus genomes or were less than 50 aa in length. After exclusions, we counted nonsynonymous divergent sites and calculated pairwise *dN/dS* of X^SR^ vs X^ST^ for 2,642 X-linked genes and for *T. dalmanni* vs Td2 for 9,525 genes using the SNAP utility (Korber et al. 2000).

To investigate the evolution of genome defense genes in the *T. dalmanni* genome, we identified *T. dalmanni* ortholog(s) and new paralogs for 80 out of 87 genes reported to be involved in RNA interference in Drosophila (Palmer et al. 2018, Tables S1 and S2). Orthologs were identified first by name from the gff annotation described above, and if absent from the annotation, a reciprocal best hit tBLASTn was used to identify matches between unannotated genes and the Drosophila piRNA gene. We fit a series of linear models to evaluate the effect of X chromosome linkage, tissue expression pattern, and status either as a gene involved in RNA interference or associated with piRNA on the rate of molecular evolution as estimated by *dN/dS*. First, *dN/dS* values were natural log transformed and passed normality testing - Shapiro-wilk test (Shapiro and Wilk 1965) was performed on 1000 randomly chosen values, P>0.10). Next 28 linear models were fit using up to three factors with and without interaction terms and using different tissue expression patterns. Models were ranked by the AICc criterion (**Table S6**) and the top model was selected (**Table S7**).

The Mcdonald-Kreitman test (**Table S8**) was performed for *T. dalmanni* orthologs of a set of genes involved in piRNA biogenesis and expression – *Argonaute-3*, *maelstrom*, *piwi*, *aubergine*, *vasa*, *hen1*, *krimp*, *armitage*, *squash*, *zucchini*, *and spindle-E*. To identify polymorphic sites within *T. dalmanni*, the Kanching ST2 library was used to represent variation among standard individuals and the Gombak and Kanching SR libraries were used to represent variation among sex-ratio individuals. Reference and alternative chromosomal pseudohaplotypes were called from the *T. dalmanni* pools using bcftools (bcftools consensus -H R and -H A). For interspecific comparisons the *T. dalmanni sp 2* (Td2) reference described above was used. GffRead was then used with the annotation to extract the alignment from each consensus and the standard McDonald-Kreitman test was performed (Egea et al. 2008).

## Supporting information

Supplementary materials list

## Data Availability

Raw data and genome assemblies used in this project are available on NCBI BioProjects PRJNA655584 (sex ratio resequencing), PRJNA391339 (*Teleopsis dalmanni s.s*. genome assembly), PRJNA662429 (multiplexed shotgun genotyping) and PRJNA659474 (*Teleopsis dalmanni sp2* aka isolate:KP12SP2M1 genome assembly). Transposable element family consensus fasta sequences for *Teleopsis dalmanni*, the MSG genotype matrix and gff3 formatted gene annotations are available on University of Maryland Digital repository at University of Maryland (DRUM), archive ID’s 1903/26380, fgxn-tuaf, and gfqi-iktk

## Acknowledgements

The authors thank Melanie Kirk, Nathaniel Lowe, Wyatt Shell, George Ru, and Gabriel Welsh for assistance with analysis, sample preparation, and fly rearing; Philip Johns and Max Brown for fly collections; Shawn Sullivan for HiC analysis; Najib El-Sayed and Suwei Zhao for HiSeq library prep and sequencing assistance; Ellen Martinson for bacterial contamination screening; and Molly Schumer, Peter Andolffato, and Wei Wang for reagents and advice on multiplexed shotgun genotyping (MSG). Funding for this work was provided by National Science Foundation grants DEB-0951816 to R.H.B., DEB-0952260 to G.S.W., by the USDA National Institute of Food and Agriculture grant 2018-67015-28199 to A.V.Z, and by the University of Maryland and the Geneseo Foundation.

## Author Contributions

JR, RB, and GW prepared the manuscript. JR, AZ, KP, CL, JW, GW, and RB analyzed the data. CH, GW provided sequencing data.

