## Supplementary materials list for "Impacts of sex ratio meiotic drive on genome structure and defense in a stalk-eyed fly"

**Josephine A Reinhardt[1], Richard H. Baker[2], Aleksey V. Zimin[3], Chloe Ladias[1], Kimberly A Paczolt[4], John Werren[5], Cheryl Hayashi[2], Gerald S. Wilkinson[4]**

1. Biology Department, State University of New York at Geneseo, Geneseo, New York, USA.
2. Sackler Institute for Comparative Genomics, American Museum of Natural History, New York, New York, USA
3. Department of Biomedical Engineering, Johns Hopkins University, Baltimore, Maryland, USA
4. Department of Biology, University of Maryland, College Park, Maryland, USA
5. University of Rochester, Rochester, New York, USA

Figure S1-S3 Legends

Figure S1 - The *T. dalmanni* X chromosome assembly and a map from backcross are largely syntenic.

Figure S2 - Tandem amplification of JASPer evidenced by spanning long read sequences.

Figure S3 - Evolution of *maelstrom* paralogs in *Teleopsis*

(Tables S1-S10 are provided in a second supplementary file (.xlsx format))

Table S1 - sequencing data used in this analysis

Table S2 - Busco results

Table S3 - Support for 6 complex inversions on the SR X compared to the ST X

Table S4 - Genes differentially expressed and varying in copy number between XSR and XST

Table S5 - TE families with differential expression or copy number between SR and ST males or candidate Y-chromosome elements

Table S6 - AICc for top 10 models predicting dN/dS between *T. dalmanni sp 1* and *T. dalmanni* sp 2

Table S7 - Fits of best linear model predicting dN/dS

Table S8 - recent adaptive evolution of piRNA genes in *T. dalmanni*

Table S9 - description of sequencing pools

Table S10 Chromosome-wide measures of variation using full or reduced dataset

**Supplementary Figure Legends**

**Figure S1** - The *T. dalmanni* X chromosome assembly and a map from backcross are largely syntenic. X chromosomal locations of markers from a multiplexed shotgun sequencing (MSG) map and a map built using short tandem repeats (STR) (Baker 2010). Both the STR map and the MSG map were built using individuals from a backcross between *T. dalmanni* from peninsular Malaysia (Gombak valley, the same location as some collections used in the present study) and Sumatra (Bukit Lawang), which shows partial reproductive isolation due to hybrid male sterilty. Most markers were colinear between the X chromosome assembly and the MSG maps, with the exception of an inverted region between 62 and 75Mbp.

**Figure S2** Tandem amplification of JASPer evidenced by spanning long read sequences. Pacific Biosciences reads were aligned to the genome region at 18.181Mbp that includes the PWWP only JASPer paralog that exhibits evidence of tandem amplification. Six reads were identified with some homology to sequences flanking both sides of the gene region. Each of the six reads contains multiple (5-6) tandem duplications of the gene region. Other reads (not shown) map to flanking sequence on either side and contain fewer tandem repeats.

**Figure S3** - Evolution of *maelstrom* paralogs in *Teleopsis*. (A) Phylogeny of *maelstrom* of selected Dipteran UNIPROT and *Teleopsis dalmanni* paralogs and orthologs using muscle for alignment and maximum likelihood phylogeny using phyML. Support values are results of approximate likelihood method (SH-like). (B) Pruned alignment of *Drosophila melanogaster maelstrom* and of *T. dalmanni maelstrom* paralogs. The shaded region indicates the *mael* functional domain. Arrows indicate positions that differ between *T. dalmanni* SR and ST males in the heterokaryotypic copies of the *Teleopsis* X-linked *mael* paralog (TelDa2_1, TelDa1_SR, TelDa1_ST).

Figure S1

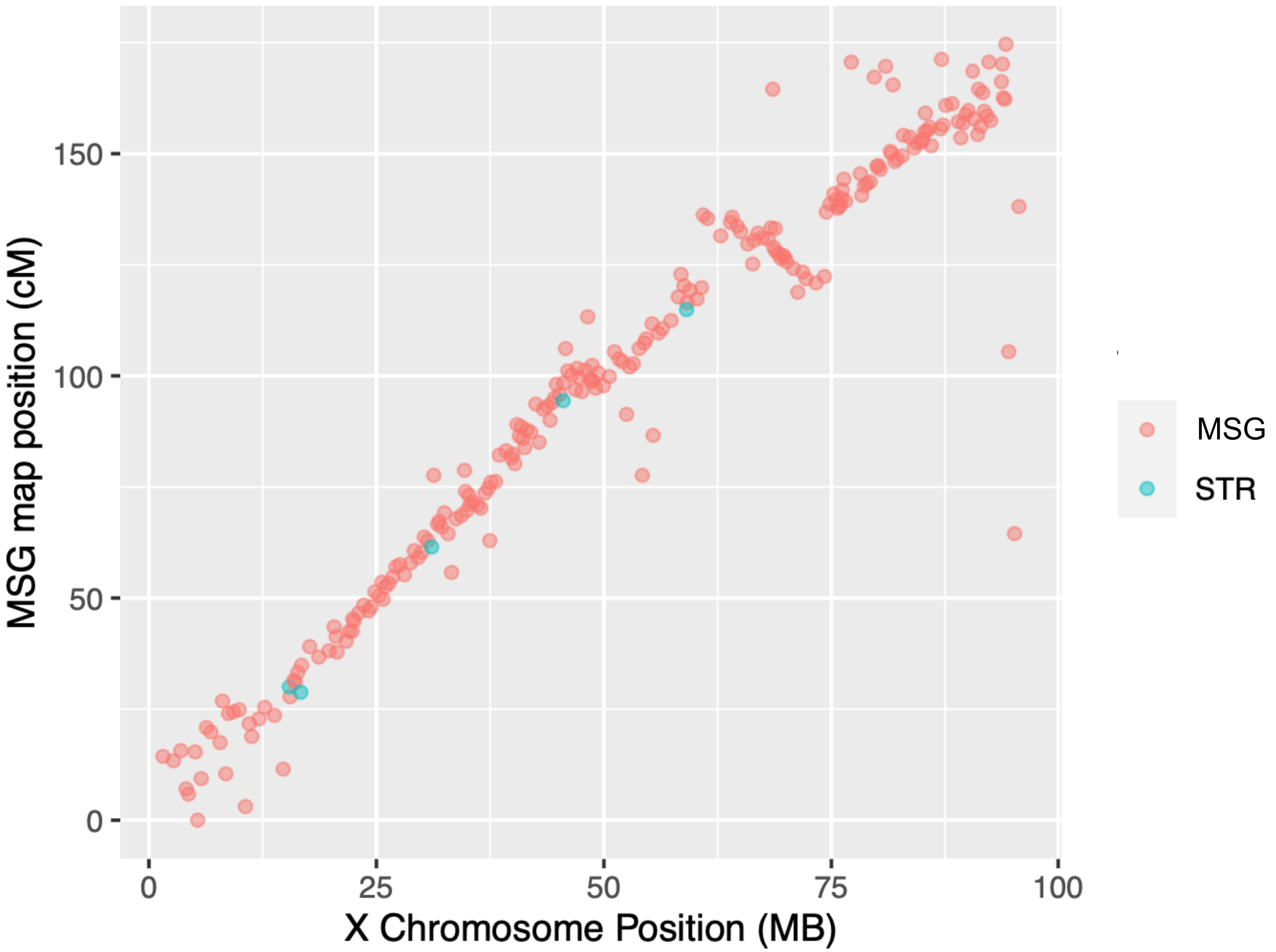

MSG map position (cM)
150
100
50
0
0
25
50
75
100
X Chromosome Position (MB)
MSG
STR

Figure S2

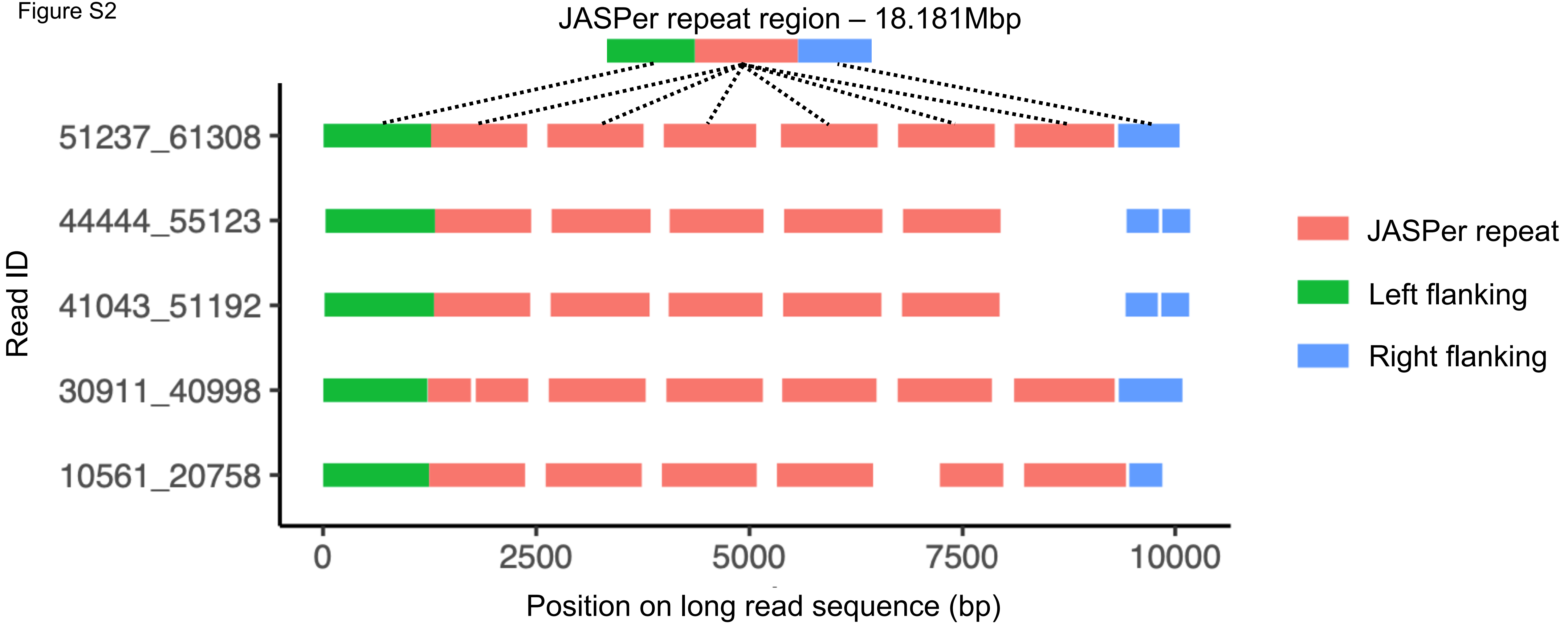
JASPer repeat region – 18.181Mbp
Read ID
51237_61308
44444_55123
41043_51192
30911_40998
10561_20758
0
2500
5000
7500
10000
Position on long read sequence (bp)
JASPer repeat
Left flanking
Right flanking

### Figure S3

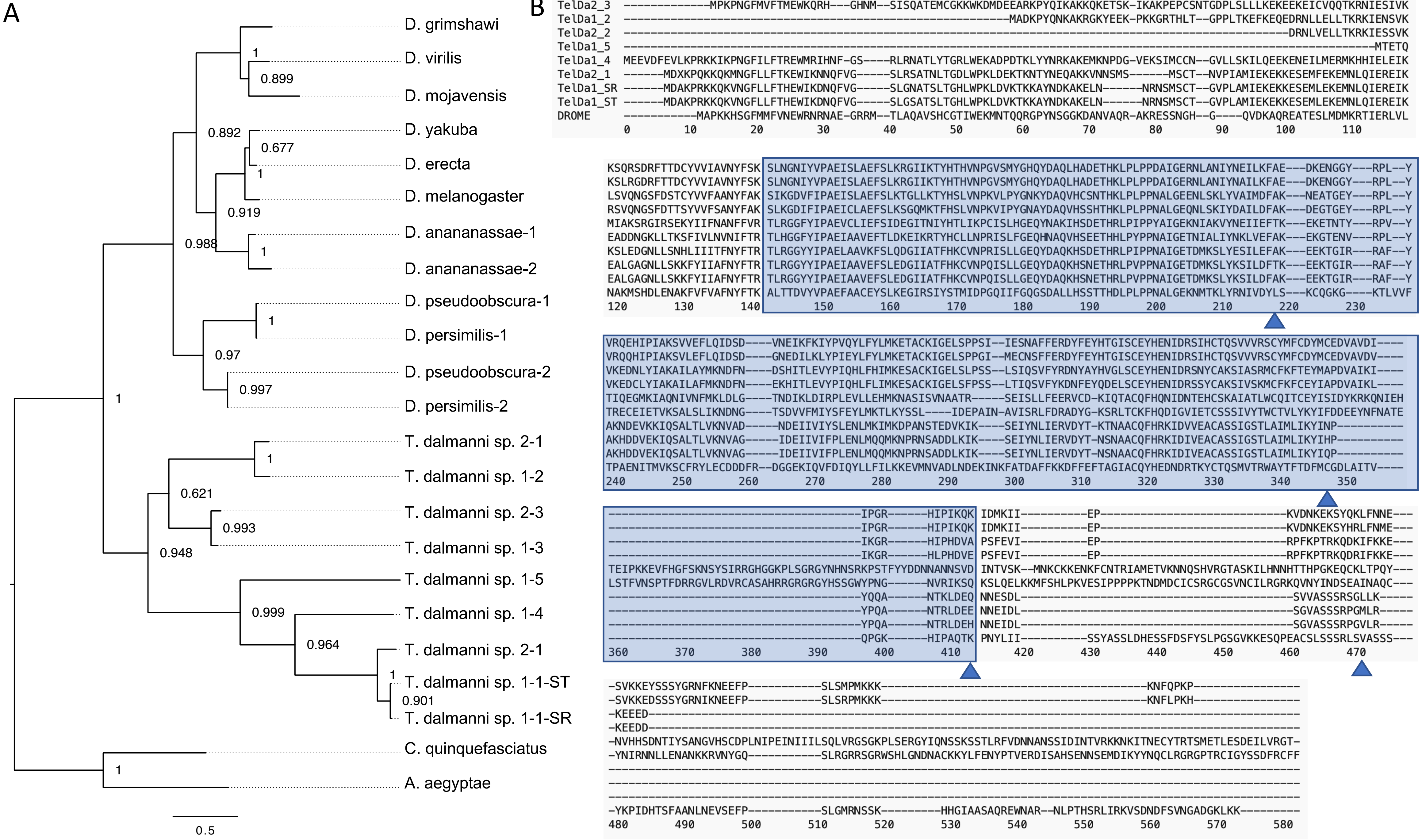
A
D. grimshawi
D. virilis
D. mojavensis
D. yakuba
D. erecta
D. melanogaster
D. ananassae-1
D. ananassae-2
D. pseudoobscura-1
D. persimilis-1
D. pseudoobscura-2
D. persimilis-2
T. dalmanni sp. 2-1
T. dalmanni sp. 1-2
T. dalmanni sp. 2-3
T. dalmanni sp. 1-3
T. dalmanni sp. 1-5
T. dalmanni sp. 1-4
T. dalmanni sp. 2-1
T. dalmanni sp. 1-1-ST
T. dalmanni sp. 1-1-SR
C. quinquefasciatus
A. aegyptae
1
0.899
0.892
0.677
1
0.919
0.988
1
1
0.97
0.997
1
1
0.621
0.993
0.948
0.999
0.964
1
0.901
1
0.5
B
TelDa1_3
TelDa2_3
TelDa1_2
TelDa2_2
TelDa1_5
TelDa1_4
TelDa2_1
TelDa1_SR
TelDa1_ST
DROME
